# The use of mutants and inhibitors to study sterol biosynthesis in plants

**DOI:** 10.1101/784272

**Authors:** Kjell De Vriese, Jacob Pollier, Alain Goossens, Tom Beeckman, Steffen Vanneste

## Abstract

Sterols are very well known for their important roles in membranes and signaling in eukaryotes. Plants stand out among eukaryotes by the large variety of sterols that they can produce, and employing them across a wide spectrum of physiological processes. Therefore, it is critical to understand the wiring of the biosynthetic pathways by which plants generate these distinct sterols, to allow manipulating them and dissect their precise physiological roles. Many enzymatic steps show a deep evolutionary conservation, while others are executed by completely different enzymes. Here, we review the complexity and variation of the biosynthetic routes of the most abundant phytosterols in the green lineage and how different enzymes in these pathways are conserved and diverged from humans,yeast and even bacteria. Based on their evolutionary conservation we discuss the use of human and yeast sterol biosynthesis inhibitors in plants, as an argument for the development of plant-tailored inhibitors of sterol biosynthesis.

## Introduction

Sterols are a class of triterpenoid lipids that consist of a hydrated phenanthrene group and a cyclopentane ring that have been a topic of great interest for researchers for many decades due to their essential physiological roles in eukaryotic organisms (Benveniste, 2004; Hartmann, 1998).

For instance, the sterol composition in membranes has a crucial impact on membrane fluidity and transmembrane export and import processes, and some sterols can even act as second messengers or signaling molecules during developmental and cellular signaling processes. The importance of sterols for eukaryotic organisms is even more apparent when looking from an evolutionary point of view, since the occurrence of sterol biosynthesis is thought to be a key evolutionary step in the advent of eukaryotic life (Galea and Brown, 2009).

Indeed, the ancient rise in atmospheric O_2_ levels to the current 21% O_2_ not only drove the evolution of the earliest eukaryotic single-cell organisms, it also allowed for the occurrence of sterol biosynthesis pathways, which require O_2_ (Galea and Brown, 2009; Mouritsen, 2005). This is contrasted by the occurrence of hopanoids in prokaryotes, which are ring-structured molecules that look similar to sterols, but that do not require O_2_ for their biosynthesis and lack a 3β-hydroxyl group, but exert analogous functions in the membranes as cholesterol (Berry *et al.*, 1993; Mangiarotti *et al.*, 2019; Saenz *et al.*, 2015). Intriguingly, the advent of sterol biosynthesis may also have acted as an early defense mechanism protecting against oxidative damage in these primitive eukaryotes, since sterols have been shown to function as a primitive cellular defense against O_2_ and reactive oxygen species (ROS) and are able to regulate cellular and organellar O_2_ entry (Galea and Brown, 2009). It is thus possible that primitive eukaryotes evolved sterols as an adaptive response to the rising atmospheric O_2_ levels, instead of just a consequence of it like previously assumed. Notably, some bacteria also produce sterols, presumably due to horizontal gene transfer (Bode *et al.*, 2003; Rivas-Marin *et al.*, 2019).

While sterols occur in all eukaryotic organisms, the types and amounts of sterols varies considerably between the different kingdoms. For instance, cholesterol is the major sterol produced in animals, whereas fungi mainly produce ergosterol. Plants, on the other hand, produce a wide variety of sterols (or phytosterols), with over 200 kinds known to date (Benveniste, 2004; Guo *et al.*, 1995; Schaller, 2004). Within the phytosterols, campesterol, stigmasterol and β-sitosterol make up the predominant molecules of the sterol profile in plants (Benveniste, 2004; Hartmann, 1998): e.g. 64% campesterol, 6% stigmasterol and 11% β-sitosterol in Arabidopsis (Benveniste, 2004; Schaeffer *et al.*, 2001). These three phytosterols have either a methyl group (campesterol) or an ethyl group (β-sitosterol and stigmasterol) on their C-24 position, and thus are also called 24-methylsterols and 24-ethylsterols, respectively (Schaller *et al.*, 1998). The balance between 24-methylsterols and 24-ethylsterols differs between plant species and is highly regulated, since their ratio has an important effect on several cellular processes (Schaller, 2003).

For instance, reproductive organs such as flowers and seedpods are negatively affected by moderate changes in the campesterol/β-sitosterol ratio, while more severe changes in the campesterol/β-sitosterol ratio have no significant effect on stem elongation (Schaller, 2003). The main function of phytosterols is the regulation of the fluidity and permeability of membranes (Schaller, 2003). They achieve this by interacting with the saturated alkyl chains of the phospho- and sphingolipids that make up the membrane bilayers, thus limiting their mobility and permeability depending on the type and amount of sterols (Hartmann, 1998). While all of the phytosterols are able to regulate membrane fluidity and permeability, their efficiency in doing so varies (Hartmann, 1998; Schuler *et al.*, 1990; Schuler *et al.*, 1991). For instance, cholesterol has the largest stabilizing effect on membranes, followed by campesterol, β-sitosterol, and stigmasterol (Grunwald, 1971; Hodzic *et al.*, 2008). Therefore, changes in the membrane sterol composition have an effect on the membrane permeability and function (Valitova *et al.*, 2010). While phytosterols are mainly present in the PM, small amounts of them have also been found in membranes of the ER (Hartmann and Benveniste, 1987), mitochondria (Meance *et al.*, 1976), vacuole (Yoshida and Uemura, 1986) and chloroplasts (Hartmann and Benveniste, 1987). Another function in membranes to which phytosterols contribute is the formation of so-called “lipid rafts”. These lipid rafts are small, dynamic membrane domains rich in phytosterols and sphingolipids, in which certain enzymes and signaling complexes are gathered (Laloi *et al.*, 2007; Malinsky *et al.*, 2013; Simon-Plas *et al.*, 2011; Simons and van Meer, 1988). Lipid rafts have been successfully identified and isolated in several plant species and detailed analyses of their composition confirmed the presence of the main phytosterols campesterol, β-sitosterol and stigmasterol, as well as other sterols, sterol glycosides and sphingolipids (Cacas *et al.*, 2012; Mongrand *et al.*, 2004; Simon-Plas *et al.*, 2011). Consequently, the phytosterol content of membranes indirectly affects enzyme activity, signal transduction, ion transport, and protein-protein and protein-lipid interactions that take place in and over these membranes (Grandmougin-Ferjani *et al.*, 1997; Schaller, 2003).

This is evidenced by the wide range of severe phenotypes that were reported for mutants defective in sterol biosynthesis. Phenotypes of such mutants include extreme dwarfism and disturbances in embryogenesis, vascularization, fertility, cell differentiation and proliferation, depending on the sterol biosynthesis step that is disturbed (Azpiroz *et al.*, 1998; Catterou *et al.*, 2001; Clouse, 2000; Guo *et al.*, 1995; He *et al.*, 2000; Piironen *et al.*, 2000; Schaller, 2003). Currently, the origin of these sterol mutant phenotypes is poorly understood. Some can be explained by defects in auxin transport (Men *et al.*, 2008; Pan *et al.*, 2009; Titapiwatanakun *et al.*, 2009; Willemsen *et al.*, 2003; Yang *et al.*, 2013) or ethylene signaling (Souter *et al.*, 2002), whereas others derive from defects in brassinosteroid signaling as campesterol serves as a biosynthetic precursor of the brassinosteroid brassinolide. Furthermore, there are indications that phytosterols can act as signaling/regulatory molecules during plant growth and development (Fujioka and Sakurai, 1997; Guo *et al.*, 1995; Lindsey *et al.*, 2003; Vriet *et al.*, 2013).

In conclusion, phytosterols not only are vital structural components of membranes, they also play key roles during plant growth and development. Therefore, the large variety of plant sterols allows plants to adapt to constantly changing environmental conditions.

## Conservation and divergence in the early sterol biosynthesis pathway

The initial pathway from which all triterpenes (including phytosterols, lanosterol and cholesterol) are derived is called the mevalonate (MVA) pathway, which is largely conserved across eukaryotes and archaea (Buhaescu and Izzedine, 2007; Lombard and Moreira, 2011) (Fig. 1). The end products of the MVA pathway are isopentenyl pyrophosphate (IPP) and dimethylallyl pyrophosphate (DMAPP), which form the primary building blocks of all isoprenoids (Goldstein and Brown, 1990).

**Fig. 1.**
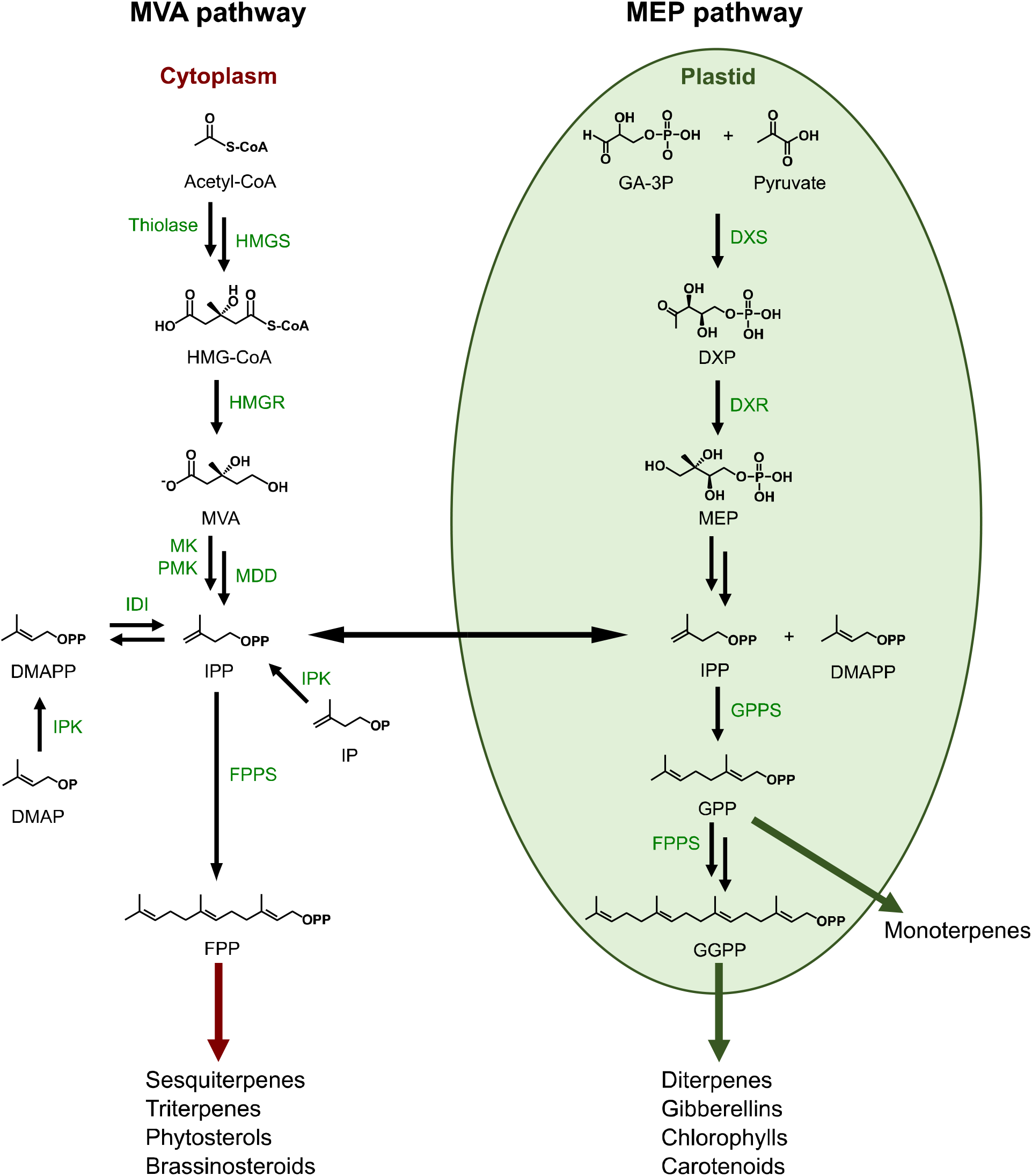
Schematic overview of the MVA and MEP pathways in *Arabidopsis thaliana*. CoA, Coenzyme A; DMAP, dimethylallyl phosphate; DMAPP, dimethylallyl pyrophosphate; DXP, 1-deoxy-D-xylulose 5-phosphate; DXR, DXP reductoisomerase; DXS, DXP synthase; FPP, farnesyl pyrophosphate; FPPS, farnesyl pyrophosphate synthase; GA-3P, glyceraldehyde 3-phosphate; GGPP, geranylgeranyl pyrophosphate; GPP, geranyl pyrophosphate; GPPS, geranyl pyrophosphate synthase; HMG, 3-Hydroxy-3-methylglutaryl; HMGR, HMG-CoA reductase; HMGS, HMG-CoA synthase; IDI, IPP isomerase; IPK, isopentenyl phosphate kinase; IP, isopentenyl phosphate; IPP, isopentenyl pyrophosphate; MDD, mevalonate 5-diphosphate decarboxylase; MEP, methylerythritol phosphate; MK, mevalonate-5-kinase; MVA, mevalonate; PMK, phosphomevalonate kinase.

The MVA pathway starts with the condensation of two acetyl-CoA molecules into acetoacetyl-CoA by acetoacetyl-CoA thiolase. An additional condensation in the next step catalyzed by HMG-CoA synthase (HMGS) results in the formation of 3-hydroxy-3-methylglutaryl-CoA (HMG-CoA). Subsequent reduction of HMG-CoA by HMG-CoA reductase (HMGR) leads to the production of mevalonate. In contrast to humans, plants often have multiple HMGR isoforms in their genomes. For instance, the *Arabidopsis* genome contains two HMGR genes that encode for three HMGR isoforms, of which HMG1 is most abundantly expressed (Enjuto *et al.*, 1994; Enjuto *et al.*, 1995). Consistent with its biochemical role in the mevalonate pathway, the pleiotropic *hmg1* phenotype can be rescued by exogenous application of squalene (Suzuki *et al.*, 2004).

In the last steps of the eukaryotic MVA pathway, MVA undergoes two phosphorylations at its 5-OH position (catalyzed by mevalonate-5-kinase (MK) and phosphomevalonate kinase (PMK)), followed by a decarboxylation (catalyzed by mevalonate 5-diphosphate decarboxylase (MDD)), resulting in IPP. This IPP, together with its derivative DMAPP that is synthesized by IPP isomerase (IDI), form the starting molecules of the pathways leading to the production of a large variety of isoprenoids (Goldstein and Brown, 1990). Archaea use a modified MVA pathway in comparison to eukaryotes, in which the last three enzymes have been replaced by other enzymes (Boucher *et al.*, 2004).

Interestingly, unlike animals and fungi, plants also have the ability to produce IPP and DMAPP via an alternative pathway: the methylerythritol phosphate (MEP) or non-mevalonate pathway (Banerjee and Sharkey, 2014; Chappell, 2002), which takes place in the plastids and is mostly used for the biosynthesis of various mono-, di- and tetraterpenoids (Laule *et al.*, 2003; Zhao *et al.*, 2013) (Fig. 1). The MEP pathway is the main pathway for IPP and DMAPP biosynthesis in bacteria, with some exceptions (Lombard and Moreira, 2011), and is obtained by plants during the endosymbiosis event with cyanobacteria that originated the plastids (Lange et al., 2000).

On the other hand, the IPP and DMAPP produced by the cytosolic MVA pathway are mainly used for the production of phytosterols, triterpenoids and sesquiterpenoids. Interestingly, many green algae species do not possess the MVA pathway and are solely reliant on the MEP pathway for isoprenoid biosynthesis (Lohr *et al.*, 2012). Notably, there are indications of crosstalk between the cytosolic MVA and plastidial MEP pathways in plants (Mendoza-Poudereux *et al.*, 2015; Tansey and Shechter, 2001). Furthermore, it was recently shown that plants express a functional homolog of the isopentenyl phosphate kinase (IPK) that was originally identified in archaebacteria as part of their modified MVA pathway (Dellas and Noel, 2010; Henry *et al.*, 2015). This enzyme catalyzes the phosphorylation of isopentenyl phosphate (IP) and dimethylallyl phosphate (DMAP) into IPP and DMAPP, respectively, thus increasing their availability for terpenoid production (Henry *et al.*, 2015; Henry *et al.*, 2018). Interestingly, IPP can be dephosphorylated back to IP by a subset of Nudix superfamily hydrolases (Henry et al., 2018). Together, these findings illustrate the highly complex metabolic regulation of IPP and DMAPP levels for terpenoid biosynthesis in plants.

Subsequently, in the cytosol, farnesyl pyrophosphate (FPP) is formed by two sequential condensation reactions, in which two IPP molecules are added to DMAPP. These condensation reactions are catalyzed by farnesyl pyrophosphate synthase (FPPS) (Kulkarni *et al.*, 2013). In the plastidial MEP pathway, on the other hand, FPP is synthesized in two steps, in which IPP and DMAPP are first converted to GPP by GPPS followed by the formation of FPP from GPP and IPP by a plastidial FPPS (Manzano *et al.*, 2016). In the cytosol, FPP can either enter the sesquiterpene biosynthesis pathway, or be further converted to squalene, a C-30 molecule which is a condensation of two FPP units catalyzed by squalene synthase (SQS) (Tansey and Shechter, 2001). Squalene is produced via this pathway in both pro- and eukaryotes, where it is the universal precursor of hopanoids and steroids, respectively. In plants, animals and fungi, squalene is further converted to 2,3-oxidosqualene by squalene epoxidase (SQE; see further) (Thimmappa *et al.*, 2014).

## Early Phytosterol biosynthesis – 2,3-oxidosqualene as a precursor for phytosterols

As mentioned before, the biosynthesis of sterols in eukaryotes begins with the epoxidation of squalene into 2,3-oxidosqualene by SQUALENE EPOXIDASEs (SQE) (Thimmappa *et al.*, 2014). Of three functional SQEs in *Arabidopsis* that can rescue SQE deficient yeast (Laranjeira *et al.*, 2015; Rasbery *et al.*, 2007), SQE1 seems to play the most predominant function, as a single mutant displays pleiotropic phenotypes in the root and shoot (Pose *et al.*, 2009; Rasbery *et al.*, 2007). However, these phenotypes were not due to the reduced sterol content of the mutant, but rather due to its hyperaccumulation of squalene (Doblas *et al.*, 2013). Moreover, the *sqe1* phenotypes could be explained by misregulation of ROS production (Pose *et al.*, 2009), unlike later sterol biosynthetic mutants that display misregulated ethylene production and auxin transport (See further) (Souter *et al.*, 2002; Souter *et al.*, 2004). This observation gives further credibility to the hypothesis that sterol biosynthesis may have evolved as an adaptation to oxidative stress (Galea and Brown, 2009). Furthermore, these data provide evidence for a primordial role for a conserved oxidosqualene biosynthesis pathway acting as the earliest section of the phytosterol biosynthesis pathway. However, the absence of completely predictable reductions in the total phytosterol levels upon interference with enzymes involved in oxidosqualene biosynthesis indicates an important gap in our understanding of how early phytosterol biosynthesis is regulated. Indeed, recently, an alternative SQE has been identified in the diatom *Phaeodactylum tricornutum*, that belongs to the fatty acid hydroxylase superfamily instead of to the flavoprotein monooxygenases like the conventional SQEs (Pollier *et al.*, 2019). This suggests that different enzymatic reactions in plant phytosterol biosynthesis can be mediated by a wider palette of enzymes than would be expected based on sequence homology to yeast and human sterol biosynthetic genes.

Depending on the plant species, there are multiple cyclization pathways that convert 2,3-oxidosqualene into different cyclic triterpene derivatives, based on the oxidosqualene cyclases (OSCs) that are present. These OSCs evolved from bacterial squalene/hopane synthases, and include cycloartenol synthase (CAS), lanosterol synthase (LAS), thalianol synthase (THAS) and β-amyrin synthase (bAS) (Sawai *et al.*, 2006; Thimmappa *et al.*, 2014). The most prominent of these pathways starts with the cyclization of 2,3-oxidosqualene into cycloartenol, which is catalyzed by the enzyme cycloartenol synthase 1 (CAS1) in *Arabidopsis* (Gas-Pascual *et al.*, 2014; Rees *et al.*, 1969; Thimmappa *et al.*, 2014). This pathway mainly produces the three major phytosterols as end-products, namely campesterol, β-sitosterol and stigmasterol, via a complex series of enzyme-catalyzed conversions. Interestingly, the bacterium *Stigmatella aurantiaca* also produces cycloartenol via a CAS enzyme that is similar to that of plants (Bode *et al.*, 2003), and a squalene monooxygenase and an OSC were found to be essential for lanosterol biosynthesis in the bacterium *Gemmata obscuriglobus* (Rivas-Marin et al., 2019).

Cycloartenol is first converted to 24-methylenecycloartenol by the addition of a methyl-group at the C-24 position by C-24 sterol methyltransferase 1 (SMT1), which is a key regulatory step of phytosterol biosynthesis (Neelakandan *et al.*, 2009; Shi *et al.*, 1996). In the next step, removal of a methyl group from the C-4 position of 24-methylenecycloartenol leads to in cycloeucalenol. In Arabidopsis, this step is catalyzed by three members of the sterol-4α-methyl oxidase 1 (SMO1) enzyme family (Darnet and Rahier, 2004). The opening of the cyclopropane ring of cycloeucalenol by cycloeucalenol cycloisomerase (CPI1) subsequently leads to the production of obtusifoliol (Benveniste, 2002). Obtusifoliol then undergoes demethylation of its C-14 position, which results in the formation of 4α-methyl-5α-ergosta-8,14,24(28)-trien-3β-ol (Rahier and Taton, 1986). This reaction is catalyzed by obtusifoliol 14α-demethylase (CYP51G1 in Arabidopsis), a cytochrome P450 enzyme. Next, 4α-methyl-5α-ergosta-8,14,24(28)-trien-3β-ol is converted to 4α-methylfecosterol by the sterol C-14 reductase FACKEL (FK). In the following step, the C-7 double bond of 4α-methylfecosterol undergoes a reduction catalyzed by the C-8,7 sterol isomerase HYDRA1 (HYD1) (Souter *et al.*, 2002), which leads to the formation of 24-methylenelophenol.

This part of the phytosterol pathway in Arabidopsis is encoded by single genes and the corresponding mutants often display strong phenotypes, including *cpi, smt1*, *cyp51*, *hyd1* and *fk*. The *cpi* mutant is characterized by increased levels of cycloeucalenol and its derivatives, and has severe defects in its growth and development (Men *et al.*, 2008). The *smt1, hyd1* and *fk* mutants have reduced phytosterol and BR levels, and are severely impaired in embryogenesis, cell polarity, root growth, gravitropism and vascular development (Diener *et al.*, 2000; Schrick *et al.*, 2000; Souter *et al.*, 2002; Topping *et al.*, 1997; Willemsen *et al.*, 2003). The abnormal vascular development phenotype of the *hyd1* and *fk* mutants could be partially rescued by crossing these mutants with auxin-resistant mutants, indicating that the *hyd1* and *fk* mutants may have disturbed auxin signaling or transport (Souter *et al.*, 2002). Similarly, the abnormal root phenotype in *hyd1* and *fk* could be rescued by crossing these mutants with a dominant ethylene-resistant mutant, suggesting they also have disturbed ethylene signaling (Souter *et al.*, 2002). Recently, tissue-specific complementation of the *hyd1* mutant suggests that many of its phenotypes can be explained by defective patterning of PIN auxin transporters, and associated defects in auxin transport (Diener *et al.*, 2000). Interestingly, the *smt1* mutant is hypersensitive to Ca^2+^ ions, since lowering the Ca^2+^ concentration in the growth medium of this mutant resulted in improved root growth, probably due to alterations in membrane permeability (Diener *et al.*, 2000).

## Phytosterol biosynthesis – parallel branches for stigmasterol and campesterol

From 24-methylenelophenol onwards, the pathway bifurcates via two separate branches, eventually resulting in either 24-ethylsterols (β-sitosterol and stigmasterol) or 24-methylsterols (campesterol) as end-products, respectively. Campesterol can subsequently be used as a precursor for brassinosteroid biosynthesis.

The 24-ethylsterol branch pathway begins with a second methylation of the C-24 position of 24-methylenelophenol by the enzymes C-24 sterol methyltransferase 2/cotyledon vascular pattern 1 (SMT2/CVP1) and C-24 sterol methyltransferase 3 (SMT3), which results in 24-ethylidenelophenol (Bouvier-Nave *et al.*, 1998; Carland *et al.*, 2010). Like with SMT1, the reaction catalyzed by SMT2/CVP1 is an important regulatory step in sterol biosynthesis, since it determines the ratio of 24-methyl- and 24-ethylsterols, which affects several developmental processes in plants (Bouvier-Nave *et al.*, 1997; Carland *et al.*, 2002; Schaeffer *et al.*, 2001). Interestingly, it is thought that SMT2/CVP1 is also able to catalyze the primary C-24 methylation catalyzed by SMT1, albeit to a lesser extend (Schaeffer *et al.*, 2001). A following double demethylation of 24-ethylidenelophenol by sterol-4α-methyl oxidase 2 (SMO2) results in the formation of Δ^7^-avenasterol (Darnet and Rahier, 2004), which is subsequently converted to 5-dehydroavenasterol by the Δ^7^-sterol-C5-desaturase DWARF7/STEROL1 (DWF7/STE1) (Choe *et al.*, 1999b; Gachotte *et al.*, 1996). Next, the sterol Δ^7^-reductase DWARF5 (DWF5) converts this 5-dehydroavenasterol into isofucosterol (Choe *et al.*, 2000). Finally, a C-24 reduction of isofucosterol by the Δ^24^-sterol-Δ^24^-reductase DIMINUTO/DWARF1 (DIM/DWF1) leads to the generation of β-sitosterol (Choe *et al.*, 1999a), which can then undergo a further C-22 desaturation by the C-22 sterol desaturase CYP710A1, resulting in the end-product of this pathway: stigmasterol (Morikawa *et al.*, 2006). However, not many details are known about this desaturation reaction in higher plants. Interestingly, in Arabidopsis, a second CYP710 enzyme (CYP710A2) is also able to produce stigmasterol from β-sitosterol, and can also produce brassicasterol from 24-epi-campesterol (Benveniste, 2002; Morikawa *et al.*, 2006).

The 24-methylsterol branch pathway starting from 24-methylenelophenol that eventually leads to the production of campesterol is similar to the first one and mostly uses the same enzymes. However, instead of first being methylated at the C-24 position by SMT2/CVP1 and SMT3 during the first step of this branched pathway, 24-methylenelophenol is directly demethylated by SMO2. This causes 24-methylenelophenol to be converted to episterol (Darnet and Rahier, 2004). The rest of the pathway consists of the same steps as the first branched pathway. First STE1 causes a desaturation of the C-5 position of episterol, which results in 5-dehydroepisterol. This is followed by a reduction of its C-7 position by DWF5, leading to 24-methylenecholesterol (Choe *et al.*, 2000). Finally, a reduction of the C-24 double bond of 24-methylenecholesterol by DIM/DWF1 yields the end-product of this pathway: campesterol. Besides its function as a structural phytosterol in membranes, campesterol also acts as a precursor for the brassinosteroid biosynthesis pathway (Choe *et al.*, 1999a; Clouse, 2011). For details about brassinosteroid biosynthesis, we refer to dedicated reviews (Choe *et al.*, 1999a; Clouse, 2011).

The *smt2*/*cvp1* mutant has increased campesterol levels and reduced β-sitosterol levels, and is characterized by moderate developmental defects, such as disturbed venation patterns in its cotyledons, serrated floral organs and a reduced stature (Carland *et al.*, 2010; Carland *et al.*, 2002). Unlike the early sterol biosynthesis mutants *smt1*, *hyd1* and *fk*, more downstream sterol biosynthesis mutants such as *smt2*/*cvp1*, *dim/dwf1*, *dwf5* and *dwf7/ste1* show no defects in embryogenesis. The *smt2*/*cvp1* mutant is smaller than the wild type, but it doesn’t demonstrate extreme dwarfism (Carland *et al.*, 2002). Although *dim/dwf1*, *dwf5* and *dwf7/ste1* affect successive steps in the conversion of episterol to campesterol, and Δ^7^-avenasterol to β-sitosterol (Choe *et al.*, 1999a; Choe *et al.*, 1999b; Clouse, 2002), the phenotypes of these mutants resemble those of brassinosteroid-deficient mutants, reflecting the importance of campesterol as a precursor of the most biologically active brassinosteroid, brassinolide. However, while these mutants are significantly smaller than wild-type plants, they don’t display the extreme dwarfism that is typical of BR biosynthesis mutants. Furthermore, the sterol profile of these mutants is vastly disturbed, with *dwf7/ste1* being almost completely devoid of campesterol (Choe *et al.*, 1999b; Choe *et al.*, 2000). These macroscopic phenotypes can be partially rescued by external application of BRs (Choe *et al.*, 1999a; Choe *et al.*, 1999b; Choe *et al.*, 2000; Klahre *et al.*, 1998; Schaller, 2003), demonstrating that they are largely caused by an impairment in downstream BR synthesis, rather than a direct effect of campesterol deficiency. However, since DIM/DWF1, DWF5 and DWF7/STE1 also catalyze the conversion steps of Δ^7^-avenasterol to β-sitosterol (Fig. 2), their respective mutants are not only deficient in campesterol, but also in β-sitosterol and stigmasterol, suggesting that the resulting defects in membrane integrity are at least partially responsible for the observed phenotypes of these mutants. This is presumably the case for the observed fertility defects, since BR application does not restore fertility in these mutants, suggesting that phytosterols play an important role during the plant reproduction that is independent from BRs (Schaller, 2004).

**Fig. 2.**
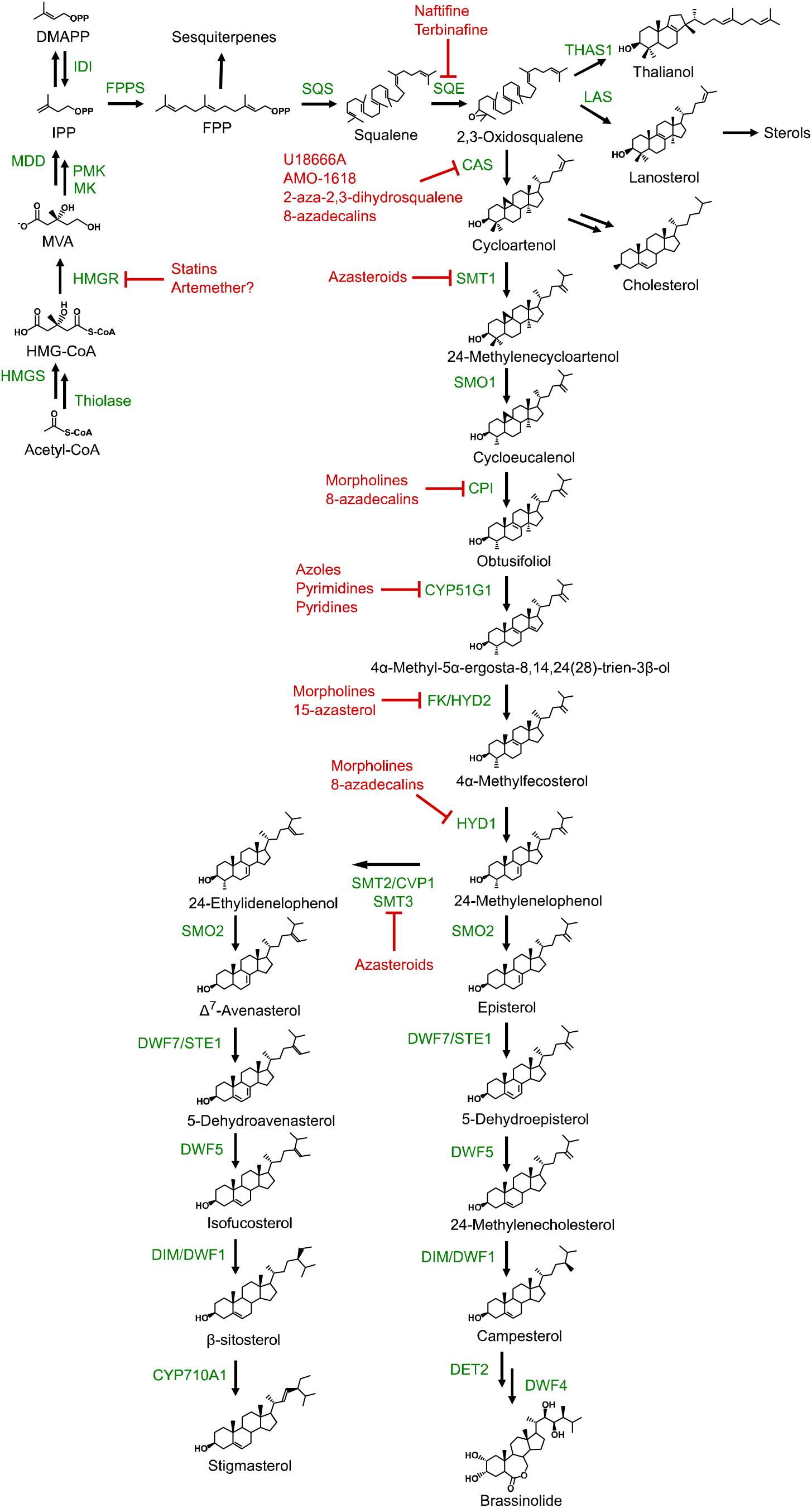
Schematic overview of the main sterol biosynthesis pathway in *Arabidopsis thaliana* and putative target sites of inhibitors. CAS, cycloartenol synthase; CoA, Coenzyme A; CPI, cycloeucalenol cycloisomerase; CVP1, cotyledon vascular pattern 1; CYP51G1, cytochrome P450 51G1; CYP710A1, cytochrome P450 710A1; DET2, DEETIOLATED2; DIM, DIMINUTO; DMAPP, dimethylallyl pyrophosphate; DWF1/5/7, DWARF1/5/7; FK, FACKEL; FPP, farnesyl pyrophosphate; FPPS, farnesyl pyrophosphate synthase; HMG, 3-Hydroxy-3-methylglutaryl; HMGR, HMG-CoA reductase; HMGS, HMG-CoA synthase; HYD, HYDRA; IDI, IPP isomerase; IPP, isopentenyl pyrophosphate; LAS, lanosterol synthase; MDD, mevalonate 5-diphosphate decarboxylase; MK, mevalonate-5-kinase; MVA, mevalonate; PMK, phosphomevalonate kinase; SMO, sterol-4α-methyl oxidase; SMT1/2/3, C-24 sterol methyltransferase 1/2/3; SQE, squalene epoxidase; SQS, squalene synthase; STE1, STEROL1; THAS1, thalianol synthase.

Furthermore, unlike *dim/dwf1*, *dwf5* and *dwf7/ste1*, the phenotypes of *smt2*/*cvp1* and the early sterol biosynthesis mutants *smt1*, *cpi*, *fk* and *hyd1* cannot be rescued by BR treatment (Carland *et al.*, 2002; Diener *et al.*, 2000; Schrick *et al.*, 2000). Since these phenotypes are independent from the downstream BR pathway, it is possible that early synthesized sterols (sterol biosynthesis intermediates) can act as signaling molecules themselves, similar to what has been shown for cholesterol in animals (Edwards and Ericsson, 1999; Farese and Herz, 1998; Vriet *et al.*, 2013). For example, accumulation of the sterol biosynthesis intermediate 4-carboxy-4-methyl-24-methylenecycloartanol (CMMC), which accumulated in a mutant defective in tethering the sterol C4-demethylation complex, interferes with auxin transport (Edwards and Ericsson, 1999; Farese and Herz, 1998; Vriet *et al.*, 2013). Also of note is that the sterol biosynthesis pathways are relatively conserved between Eukaryotes, with diatoms and yeast using mostly similar or identical enzymes as the higher plants, albeit sometimes in a different order, which explains the difference in end products obtained (e.g. ergosterol in yeast and brassicasterol/campesterol in diatoms) (Fabris *et al.*, 2014). Overall, these mutants of early and late steps of the sterol biosynthesis pathway have been excellent tools in aiding our understanding of plant sterol biosynthesis and the role of sterols in plant growth and development.

## Cholesterol biosynthesis in plants

The major sterols in plants are β-sitosterol, campesterol and stigmasterol, but many plants also produce cholesterol to some degree (Behrman and Gopalan, 2005). While the cholesterol levels in plants are usually low (100 - 1000 times lower compared to animals), cholesterol makes up a significant portion of the sterol content in some plant species (e.g. more than 10% in *Solanaceae*) (Sonawane *et al.*, 2016). Furthermore, it has been shown to serve several functions in various plant species, including as membrane component, leaf surface lipid, and precursor for several plant metabolites such as steroidal glycoalkaloids (SGAs) and phytoecdysteroids (Cardenas *et al.*, 2015; Dinan, 2001; Japelt and Jakobsen, 2013; Milner *et al.*, 2011).

Cholesterol is the major sterol in animals, in which the cholesterol biosynthesis pathway has been extensively studied and characterized (Nes, 2011), while cholesterol biosynthesis in plants is still not fully understood. Recently, several genes and enzymes involved in cholesterol biosynthesis in tomato plants were identified by analyzing transcript and protein co-expression data, as well as a combination of functional assays (Sonawane *et al.*, 2016). These data demonstrated the involvement of 12 enzymes in the tomato cholesterol biosynthesis pathway, of which several also function in the phytosterol biosynthesis pathway to catalyze highly related enzymatic conversions. Furthermore, the other enzymes that are specific for the cholesterol biosynthesis pathway seem to have evolved through gene duplication and divergence from phytosterol biosynthetic enzymes (Sonawane *et al.*, 2016). Unlike animals, cholesterol biosynthesis in plants does not seem to start from 2,3-oxidosqualene cyclization into lanosterol by LAS (Sonawane *et al.*, 2016). Instead, the OSC involved is CAS, after which cycloartenol is not only used for phytosterol biosynthesis, but also cholesterol biosynthesis (Fig. 2). Indeed, in tomato and potato plants it was shown that sterol side chain reductase 2 (SSR2) is a key enzyme in cholesterol biosynthesis that catalyzes the conversion of cycloartenol into cycloartanol, the first committed step in cholesterol biosynthesis (Sonawane *et al.*, 2016). However, while LAS probably doesn’t contribute significantly to cholesterol biosynthesis, LAS genes were identified in several plant species, including Arabidopsis (Kolesnikova *et al.*, 2006; Sawai *et al.*, 2006; Suzuki *et al.*, 2006). Furthermore, it was shown that LAS1 overexpression in Arabidopsis significantly increases the phytosterol levels while *las1* knockout mutants do not have phytosterols derived from lanosterol, indicating that there exists an alternative phytosterol biosynthesis pathway that is dependent on LAS (Ohyama *et al.*, 2009). The existence of alternative pathways contributing to phytosterol biosynthesis could explain why phytosterol levels in *cas1* mutants remain unchanged, despite a strong defect in cycloartenol synthase activity as indicated by the accumulation of 2,3-oxidosqualene (Babiychuk *et al.*, 2008).

## Chemical inhibitors of key steps in the plant sterol biosynthesis pathway

Besides mutants, another way in which sterol biosynthesis can be disrupted is through the action of chemical inhibitors that target specific steps of the sterol biosynthesis pathway (Fig. 2, Fig. 3). Indeed, sterol biosynthesis inhibitors have proven to be effective tools to probe and investigate sterol biosynthesis pathways across the different kingdoms. Many of the currently used sterol biosynthesis inhibitors have seen commercial use as fungicides and antimycotic drugs, and some can even be used to regulate plant growth (Lenton, 1987; Leroux *et al.*, 2008). Since the sterol biosynthesis pathways of plants, animals and yeast share many similar conversion steps that are catalyzed by semi-conserved enzymes, several of the most used sterol biosynthesis inhibitors function across kingdoms (Ator *et al.*, 1992). Nevertheless, there still exist clear differences in the sterol biosynthesis pathways between the kingdoms, leading to different sensitivities and specificities of sterol biosynthesis inhibitors (Nes, 2011). The following paragraphs will go into more detail about some of the most active and most used sterol biosynthesis inhibitors in plants, and their presumed targets. The compounds discussed and their presumed targets in Arabidopsis are indicated in Fig. 2. The numbers in brackets behind the discussed compounds correlate to their numbers in Fig. 3.

**Fig. 3.**
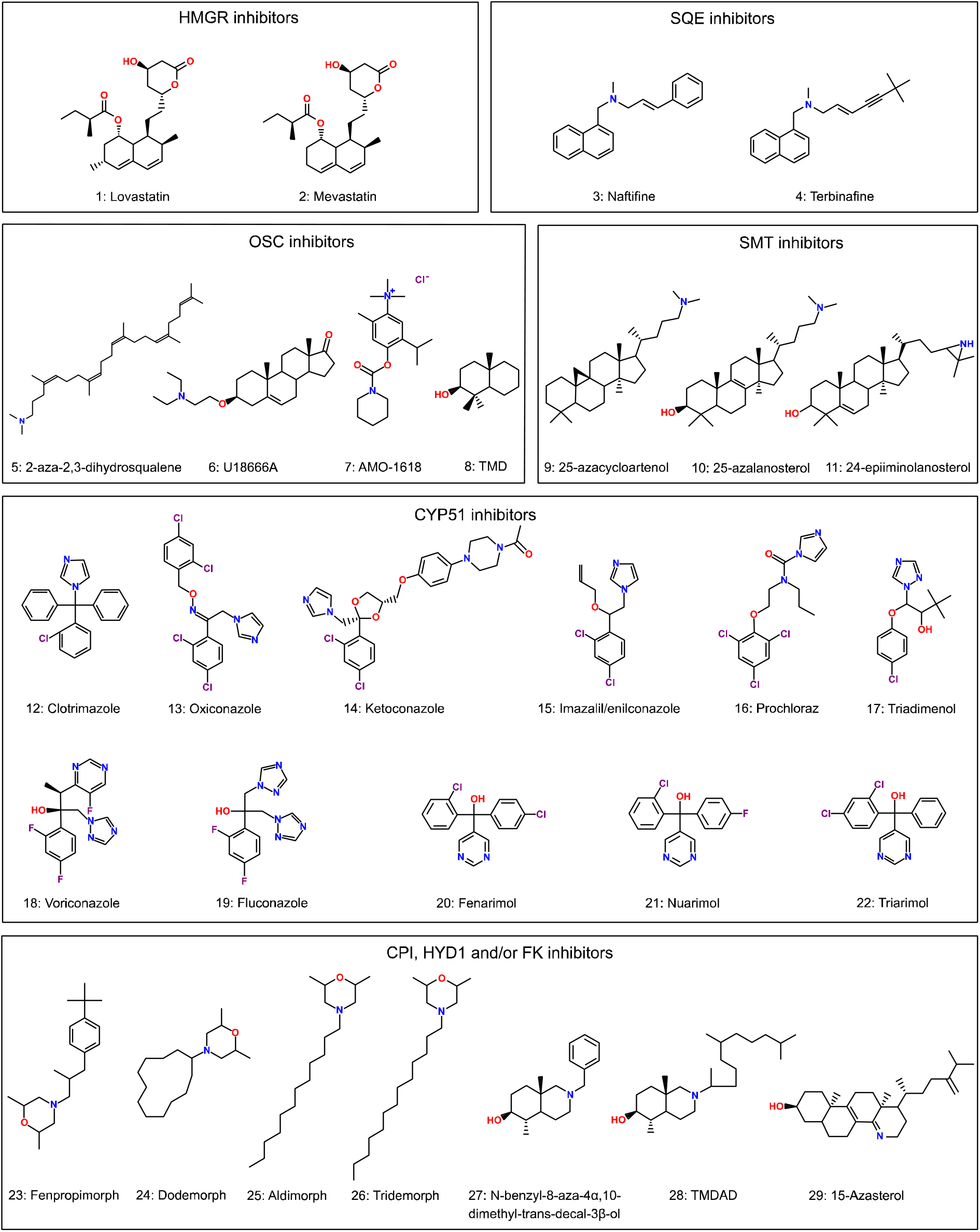
Structures of sterol biosynthesis inhibitors organized according to their putative targets.

Statins potently inhibit human HMGR activity by occupying the HMG-CoA binding site (Istvan and Deisenhofer, 2001). Because HMGR is a rate-limiting enzyme in MVA biosynthesis, statin-based medication is widely used to lower cholesterol levels (reviewed in (Davies *et al.*, 2016). In several plant species, statins, such as lovastatin **(1)** (or mevilonin) and mevastatin **(2)** (or compactin), reduce root growth and sterol biosynthesis (Bach and Lichtenthaler, 1982, 1987; Kim *et al.*, 2014; Soto *et al.*, 2011), demonstrating that statins can also be used as HMGR inhibitors in plant sterol research.

Two allylamine fungicides, namely naftifine **(3)** and terbinafine **(4)**, are potent non-competitive SQE inhibitors in fungi (Birnbaum, 1990; Ryder, 1991; Nowosielski et al., 2011). Docking analyses on modelled SQE suggest that terbinafine binding causes a conformational change that blocks one mode of substrate binding, while changing the geometry of another. (Nowosielski *et al.*, 2011). Although plant SQEs can complement yeast SQE deficient mutants (Rasbery et al., 2007), they are not highly sensitive to these inhibitors (Yates et al., 1991, 1992; Wentzinger et al., 2002). This is not surprising as single amino acid substitutions in yeast SQE are sufficient to confer terbinafine resistance (Leber *et al.*, 2003). Yet, the *sqe1-5* mutant is hypersensitive to terbinafine (Pose et al., 2009). On the other hand, some organisms such as the diatom *P. tricornutum* are completely insensitive to terbinafine as they use alternative SQEs (Fabris et al., 2014; Pollier et al., 2019).

Squalestatins (also called zaragozic acids), are highly potent and specific competitive inhibitors of rat SQS, with apparent subnanomolar *Ki* values (Baxter et al., 1992; Bergstrom et al., 1993). Also in plants, squalestatins are highly potent, as they inhibit SQS in BY-2 cell suspensions with an IC_50_ value of 5.5 nM, possibly via an irreversible inhibition mechanism (Hartmann et al., 2000; Wentzinger et al., 2002). Exogenous application of squalestatin activates transcriptional responses also seen in lovastatin-treated plants and impairs the plants fertility (Suzuki et al., 2004). The Arabidopsis genome encodes only a single functional SQS (SQS1; Busquets et al., 2008), but has not yet been subjected to mutant analysis.

Over the years, several compounds have been identified that inhibit OSCs to varying degrees by mimicking the carbocationic intermediates formed during the cyclization of 2,3-oxidosqualene. Some examples of OSC inhibitors that have been successfully utilized in plants are 2-aza-2,3-dihydrosqualene **(5)** (Duriatti et al., 1985; Cattel et al., 1986), U18666A **(6)** (Duriatti et al., 1985; Cattel et al., 1986) and AMO-1618 **(7)** (Douglas and Paleg, 1978, 1978, 1981). Another class of OSC inhibitors are the 8-azadecalins, such as 4,4,10β-trimethyl-trans-decal-3β-ol (TMD) **(8)** and its derivatives (Ruhl et al., 1989; Raveendranath et al., 1990; Hoshino et al., 1995). However, the 8-azadecalins also inhibit other enzymes besides OSCs (such as cyclopropyl sterol isomerase, C-14 sterol reductase and C-8,7 sterol isomerase), thus potentially leading to off-target effects.

In Arabidopsis, the C-24 sterol methyltransferase SMT1 catalyzes the transfer of a methyl group from S-adenosyl-L-methionine to cycloartenol (Benveniste, 1986; Bouvier-Nave et al., 1998; Diener et al., 2000), leading to the formation of Δ^5^ C-24 alkyl sterols. Since SMT1 only occurs in plants and fungi, and not in animals, it is an interesting target for studying phytosterol biosynthesis. SMT2/CVP1 and SMT3 are mainly responsible for a second methyl addition, thus resulting in an ethyl side-chain addition on the C-24 (Schaeffer et al., 2001; Carland et al., 2010). Therefore, the regulation of the SMT enzymes determines the sterol composition in plants. Many compounds have been designed over the years to act as SMT inhibitors (Nes, 2000). These inhibitors can be broadly classified in three groups: 1) substrate analogues that act as inactivators of the enzyme, 2) substrate analogues that resemble high-energy intermediates, and 3) product analogues. While these compounds are generally designed in fungal systems, some of them have been shown to inhibit SMT1 and SMT2/CVP1 in plants as well, including the azasteroid inhibitors 25-azacycloartenol **(9)** (Rahier et al., 1980; Schmitt et al., 1981; Rahier et al., 1986; Mangla and Nes, 2000), 25-azalanosterol **(10)** (Rahier et al., 1984) and 24-epiiminolanosterol **(11)** (Tal and Nes, 1987), which are carbocationic transition state analogues of the substrates of these enzymes (Rahier et al., 1984).

The 14α-methylsterol demethylase enzyme in plants (obtusifoliol 14α-demethylase) catalyzes the demethylation of obtusifoliol (Lepesheva and Waterman, 2007). This enzyme is a cytochrome P450 dependent monooxygenase (CYP51G1 in Arabidopsis) (Benveniste, 1986; Lepesheva and Waterman, 2007). In fungi and animals, the best studied and most widely used inhibitors of P450s are the azoles, which are a popular type of antifungal compounds that are used for both agricultural and medical purposes (Becher and Wirsel, 2012). Two subclasses of the azoles are the imidazoles, such as clotrimazole **(12)**, oxiconazole **(13)**, ketoconazole **(14)**, imazalil (enilconazole) **(15)** and prochloraz **(16)**, and the triazoles, such as triadimenol **(17)**, voriconazole **(18)** and fluconazole **(19)**. They are nitrogen-containing heterocyclic compounds that form an effective class of fungicides by non-competitively binding to the ferric ion of the heme group of fungal CYP51, thus preventing it from binding its substrate (Rogerson et al., 1977; Warrilow et al., 2013). Despite being primarily used as fungicides, several azole compounds also have an effect on plants to a varying degree, where they generally cause growth inhibition, which may be due to interference with downstream BR biosynthesis (Scheinpflug and Kuck, 1987; Vanden Bossche et al., 1987; Rozhon et al., 2013; Fabris et al., 2014). However, while nanomolar concentrations of azoles are usually sufficient to inhibit ergosterol biosynthesis in fungi, micromolar concentrations or higher are often needed to obtain a similar inhibitory effect on phytosterol biosynthesis in plants and diatoms (Vanden Bossche et al., 1987; Fabris et al., 2014). The cytochrome P450s are a superfamily of enzymes (Xu et al., 2015), that are often sensitive to imidazoles (Murray, 1999). Of note is that azoles can be found or even designed that display a certain degree of preference towards specific cytochrome P450 enzymes. Well-known examples in plants are uniconazole as an inhibitor of CYP707As that are involved in abscisic acid catabolism (Saito *et al.*, 2006), and brassinazole as an inhibitor of CYP90B1 that is involved in brassinosteroid biosynthesis (Asami *et al.*, 2000). Importantly, both uniconazole and brassinazole can inhibit CYP90B1 activities, suggesting that one should be wary of off-target side effects when using these inhibitors, especially when using them at high concentrations. Recently, analysis of crystals of CYP90B1 in complex with uniconazole and brassinazole demonstrated important differences in binding conformation (Fujiyama *et al.*, 2019) which highlighted the importance of using crystal structures of plant CYPs to guide the design of new, more-specific inhibitors.

Besides the azoles, also pyrimidine-type fungicides, such as fenarimol **(20)**, nuarimol **(21)** and triarimol **(22)**, and pyridine-type fungicides are thought to affect the CYP51 ortholog and other cytochrome P450 enzymes, such as CYP710A1 in plants (Shive and Sisler, 1976; Schmitt and Benveniste, 1979; Buchenauer and Rohner, 1981; Burden et al., 1987; Scheinpflug and Kuck, 1987; Mercer et al., 1989; Leroux et al., 2008; Oh et al., 2015). Overall, these compounds all cause strong reductions in root- and shoot growth with varying potency, and are phytotoxic at high concentrations due to a severe reduction in phytosterols and an accumulation of 14α-methylsterols (Burden et al., 1987; Lurssen, 1987).

CPI, C-14 sterol reductase and C-8,7 sterol isomerase are enzymes that catalyze similar reactions at different stages of the sterol biosynthesis pathway (Benveniste, 1986), making them shared targets for molecular inhibition. An important class of inhibitors that target these enzymes are the morpholine fungicides, such as fenpropimorph **(23)**, dodemorph **(24)**, aldimorph **(25)** and tridemorph **(26)** (Kerkenaar et al., 1981; Baloch et al., 1984; Baloch and Mercer, 1987; Mercer et al., 1989; Marcireau et al., 1990). These compounds exert their fungitoxicity by inhibiting C-8,7 sterol isomerase (nM concentrations) and/or C-14 sterol reductase (µM concentration) in fungi and yeast, with different morpholines having different specificities (Kerkenaar et al., 1981; Baloch et al., 1984; Baloch and Mercer, 1987; Kerkenaar, 1987; Marcireau et al., 1990). For instance, while fenpropimorph can effectively inhibit both the C-8,7 sterol isomerase and C-14 sterol reductase in fungi, tridemorph primarily inhibits the C-8,7 sterol isomerase (Baloch et al., 1984; Kerkenaar, 1987). However, morpholines also function in plants, albeit less potently and less specifically, where they have been shown to inhibit HYD1 (the plant C-8,7 sterol isomerase) (Rahier et al., 1986; Taton et al., 1987), FK (the plant C-14 sterol reductase) (Mercer et al., 1989; Taton et al., 1989; He et al., 2003) and CPI (Taton et al., 1987) to a varying degree. Similarly, fenpropimorph treatment caused alterations in the sterol content of the diatom *P. tricornutum* that could be explained by inhibition of multiple enzymes involved in its sterol biosynthesis pathway (Fabris et al., 2014). While fenpropimorph is the most active and commonly used morpholine in plants, it requires relatively high concentrations to function (30 - 100 μM), is unstable and relatively expensive. Plants treated with morpholines have a disturbed sterol profile and growth impairments, similar to mutants defective in the targeted enzymes (Bladocha and Benveniste, 1983; Burden et al., 1987; He et al., 2003). However, while the morpholine compounds disturb the normal sterol profile of plants, they are generally not phytotoxic (Bladocha and Benveniste, 1983; Taton et al., 1987, 1987). Interestingly, in plants, 8-azadecalins such as N-benzyl-8-aza-4α,10-dimethyl-trans-decal-3β-ol **(27)** and N-(1,5,9-trimethyldecyl)-4α,10-dimethyl-8-aza-trans-decal-3β-ol (TMDAD) **(28)** have been shown to be more potent inhibitors of HYD1 and CPI than the morpholines (Rahier et al., 1985; Taton et al., 1987). A strong, more specific inhibitor of C-14 sterol reductases is the antifungal agent 15‐aza‐24‐methylene‐D‐homocholesta‐8,14‐dien‐3β‐ol (15-azasterol) **(29)**, which has been used to inhibit FK in several plant species, including Arabidopsis (Schrick et al., 2002) and bramble cells (Schmitt et al., 1980).

While most of the abovementioned sterol biosynthesis inhibitors have been used to inhibit plant growth and study plant and diatom sterol biosynthesis to some degree, it is clear that many of these compounds originate as antifungal compounds for which the effect in plants is often not completely understood. Indeed, much of the underlying mechanisms of these inhibitors in plants and diatoms are still not completely clear and are often presumed based on their function in fungi and/or animals. It should also be noted that only limited recent data is available for most of these inhibitors in plants, as evidenced by the relatively old sources referenced in the last paragraphs. This further supports the notion that the current knowledge and toolset of sterol biosynthesis inhibitors in plants is lacking. The identification of more active compounds that selectively target specific enzymes in the plant sterol biosynthesis pathway through a systematic approach, informed by crystal structures, would therefore be highly welcomed to study sterol biosynthesis in the green lineage.

## Acknowledgments

K.D.V. is funded by the Special Research Fund Ghent University.

## Author Contributions

"Conceptualization, K.D.V and S.V..; All authors contributed to the writing and revision of the manuscript

